# The *Neisseria meningitidis* iron acquisition protein HpuA moonlights as an adhesin and inhibits host cell migration

**DOI:** 10.1101/2022.02.25.481912

**Authors:** Gabrielle A. Shortt, Xiaoyun Ren, Brianna M. Otto, Joanna K. MacKichan

## Abstract

*Neisseria meningitidis* can cause meningococcal disease, a rapidly developing and potentially fatal infection. Despite this, it normally resides as a commensal in the nasopharynx of healthy individuals. The mechanisms by which meningococci access deeper tissues remain unknown. Epidemiological data suggest that mucosal disruptions increase the risk of meningococcal disease. We previously investigated whether meningococci inhibit host cell wound repair, enhancing invasive disease risk. Here, using genome sequencing and a collection of closely related household isolates that differ in their ability to inhibit host wound repair, we identify the responsible meningococcal factor. This protein, HpuA, has previously been characterized as part of a bipartite heme acquisition transporter. We constructed mutants to demonstrate that HpuA, but not HpuB, inhibits wound repair, acts as an adhesin for epithelial cells, and promotes cellular invasion. We showed this was not due to iron starvation resulting from the bacteria, differences in growth rate, or manipulation of host haptoglobin. Heterologous expression of HpuA in *E. coli* mediated adherence to 16HBE cells in an HpuA-dependent manner and conferred an aggregative phenotype onto *E. coli*, suggesting that HpuA may play a role in the formation of microcolonies on host cells. We also demonstrated that iron supplementation of meningococci restored the inhibition of wound repair in strains lacking HpuA (NZCM112, Δ*hpuA* mutant) to levels seen with the wild type. This was also seen with unrelated carriage strains previously shown not to inhibit wound repair. Iron supplementation also increased adherence and invasion of meningococci for strains lacking HpuA, while not affecting those that expressed HpuA. These findings suggest there may be a second meningococcal protein that inhibits wound repair. Together, these results suggest that HpuA is an important meningococcal virulence factor with multiple moonlighting functions, mediating adherence, invasion, inhibition of wound repair, and bacterial aggregation.

**Author Summary:** *Neisseria meningitidis* causes meningococcal disease, a potentially fatal and rapidly developing illness that most often occurs in children. Despite this, the bacteria are frequently carried harmlessly as part of the normal airway microflora in healthy people, only rarely causing invasive disease, which involves replication in the bloodstream or central nervous system. It remains unknown precisely how the bacteria reach the deeper tissues from the airways, though some epidemiological evidence suggests that wounds or disruptions to the airways may increase risk. Here, we show that a *N. meningitidis* protein, HpuA, moonlights from its usual job of acquiring nutrients from the host, to enable the bacteria to adhere to and invade host cells, as well as inhibiting wound closure. Furthermore, we also show that meningococci that lack HpuA acquire the ability to inhibit wound repair when they are supplemented with iron, suggesting that there are additional meningococcal proteins to be discovered that may inhibit wound repair.

## Introduction

### *N. meningitidis* infection and carriage

*Neisseria meningitidis*, a Gram-negative pathogen, is highly adapted to its human host (1). The bacteria are carried asymptomatically in about 10% of the general population as part of the commensal flora of the human nasopharynx (2, 3). However, on rare occasions, the bacteria can invade into the bloodstream and deeper tissues, causing meningococcal disease, a rapidly developing and potentially devastating disease that occurs mainly in children (4).

*N. meningitidis* isolates are highly heterogeneous, with some strain and sequence types strongly associated with invasive disease, with others more likely to be present during asymptomatic carriage (5, 6). Both bacterial and host factors are known to contribute to the risk of invasive disease (7, 8). However, the overall genetic heterogeneity of *N. meningitidis* isolates has obscured the identification of virulence factors (9).

The carriage state is asymptomatic but important to understand, as it may determine outcomes such as invasive disease, long-term carriage, transmission to new hosts, or immune clearance. While understanding bacterial factors required for asymptomatic carriage is hampered by a lack of relevant animal models, a picture is emerging. During carriage, meningococci are associated with microvilli on the surface of nonciliated epithelial cells, while models that include airway mucus suggest the bacteria are generally found in the mucus and rarely interact with host cells (10, 11). The bacteria encounter extremely limited iron concentrations in the mucus and express a range of iron- and heme-acquisition proteins (12, 13). Even in the presence of mucus, however, *N. meningitidis* is still seen in close association with host cells (11), an association that may be important for iron piracy by the bacteria. *N. meningitidis* have been shown to acquire iron directly from host epithelial cells, in a process that has been shown to be TonB-dependent and that may involve degradation products of host ferritin (14, 15). While these iron sources have been suggested to not include heme, at least one heme utilization protein has been shown to be important in the interactions beween *N. meningitidis* and epithelial cells (16). In addition, intracellular *N. meningitidis* inside nasopharyngeal epithelial cells have been shown to perturb transferrin transport, which may be an alternative strategy to acquire host intracellular iron stores, similar to one deployed by *Helicobacter pylori* (17, 18).

Colonizing *N. meningitidis* bacteria also evade the antimicrobial effects of the immune response, exhibiting resistance to complement and phagocytosis, and using phase and antigenic variation to evade antibody recognition (19). Some hypervirulent *N. meningitidis* strains also specifically interfere with innate immune signaling pathways, including TNF-α and NF-κB pathways, likely dampening the ability of the host to activate an immune response (20–22). *N. meningitidis* also interacts with other commensal bacteria in the airways, including other non-pathogenic *Neisseria* species, which may have an antagonistic effect on *N. meningitidis* (23, 24)

### *N. meningitidis* and epithelial disruptions or wounds

The means by which *N. meningitidis* traverse the nasopharyngeal mucosa to cause invasive disease remain unclear (25). Active translocation following bacterial internalization is hypothesized, although fully differentiated epithelial cells may prevent dissemination of the bacteria to the deeper tissues (17, 26–28). In some cases, disruptions to the nasopharyngeal epithelium, such as viral infections, increase the risk of meningococcal disease (29, 30). Climate conditions may also increase the risk of meningococcal disease; prior to the deployment of the MenAfriVac vaccine, large outbreaks of meningococcal disease in the African “meningitis belt” were strongly associated with dust and particulate-filled wind known as the Harmattan (31–33). Elsewhere, smoking, first- or secondhand, can also increase the risk of invasive disease (34). We therefore previously investigated whether some meningococcal isolates could slow the repair of epithelial wounds, thus prolonging the opportunity for the bacteria to gain access to the bloodstream (35). An in vitro wound repair assay revealed that disease-associated meningococcal isolates inhibited cell migration, whereas carriage-associated isolates were highly variable (35). Some carriage-associated isolates inhibited cell migration to a similar degree as the invasive strains, while others had no effect on wound repair. Our previous study ruled out a role for many different meningococcal surface structures and virulence factors, e.g., the capsule or Type IV pili, in the inhibition of wound repair (35). However, the high level of genetic heterogeneity made it difficult to identify the bacterial factor mediating the inhibition of wound repair.

In the current study, we used closely related isolates that differed in their ability to inhibit wound repair to identify the factor responsible. This group of isolates included one collected from a patient and two similar nasopharyngeal carriage isolates from the patient’s healthy household contacts. The three isolates were indistinguishable by conventional laboratory typing methods, but one of the carriage isolates had lost the ability to inhibit wound repair. Comparison of the genome sequences from the three isolates revealed only one variation predicted to affect protein expression in the isolate that did not inhibit wound repair. This variation was a frameshift mutation resulting in a truncated version of HpuA, a haptoglobin:haemoglobin binding protein. Targeted mutagenesis of *hpuA* in the disease isolate abrogated its ability to inhibit wound repair, while targeted mutagenesis of the co-transcribed gene directly downstream of *hpuA*, *hpuB*, had no effect. We subsequently found that the *hpuA* mutant had an unexpected defect in adherence to epithelial cells, suggesting a novel moonlighting function for *N. meningitidis* HpuA. Microscopy of adherent bacteria also revealed that *N. meningitidis* expressing HpuA formed microcolonies on tissue culture cells, suggesting an aggregative function for HpuA. Heterologous expression of inducible HpuA in a protein expression strain of *E. coli* conferred the ability of those bacteria to adhere to epithelial cells and to aggregate, but only when HpuA was induced. We further showed that iron supplementation altered the interactions between *N. meningitidis* strains lacking HpuA and host epithelial cells, suggesting there may be additional *N. meningitidis* factors affecting wound repair and adherence to be discovered. These results are the first to demonstrate a moonlighting function for a *N. meningitidis* iron acquisition protein and suggest that there are likely others to be identified.

## Results

### Closely related household contact isolates differ in their ability to inhibit epithelial cell wound repair

As part of our efforts to identify the *N. meningitidis* factor responsible for the inhibition of wound repair, we screened a collection of isolates that were derived from a household contact study, carried out in Auckland, New Zealand, in the 1990s, during the peak of the serogroup B *N. meningitidis* epidemic (36). Patient isolates were cultured from blood or cerebrospinal fluid samples from the patient, while carriage isolates were collected from healthy household contacts of the patients by nasopharyngeal swabs. For this study, we focused on disease and carriage isolates that were indistinguishable by standard laboratory typing techniques, enabling us to study isolates that were likely closely related.

### Selection of meningococcal isolates for study

We chose one household contact group to study further. The isolate from the index patient (NZ97/052) belonged to the strain type C:2b:P1.2,5, and the sequence type (ST)-8 hypervirulent lineage. While this disease case occurred during the serogroup B meningococcal epidemic, it did not belong to the dominant strain type that drove the epidemic (37). Two additional carriage isolates (NZCM111 and NZCM112) of the same strain and sequence type were recovered from household contacts of the patient. An Oris cell migration assay was used to measure the inhibition of cell migration by the three *N. meningitidis* isolates. Bronchial epithelial cells were cultured in 96-well plates, with rubber stoppers preventing adherence to the center of each well. Following removal of the rubber stoppers, bacteria were added, and the degree of “wound’ closure was assessed following 16 hours. Despite the similarity of the three isolates, only NZ97/052 and NZCM111 inhibited wound closure, whereas co-incubation of NZCM112 with the epithelial cells resulted in significantly more wound closure (Fig 1A). Because wound closure is assessed following 16 hours of co-incubation, we asked whether the NZCM112 phenotype we observed could be due to impaired growth under the assay conditions. While we confirmed that similar numbers of colony forming units (CFU) of each bacterial strain were added to the Oris assay, after 16 hours of growth, we recovered fewer NZCM112 CFU, demonstrating that this strain has a growth impairment relative to NZ97/052 (Fig 1B). To see if reduced bacterial numbers were responsible for the altered ability to inhibit host cell wound repair, we infected cells with starting multiplicities of infection (MOI) of 10, 15, or 20. The starting MOI of each strain did not alter the inhibition of wound repair (Fig 1C), suggesting that the phenomenon of wound repair inhibition is independent of starting bacterial numbers.

**Figure 1.**
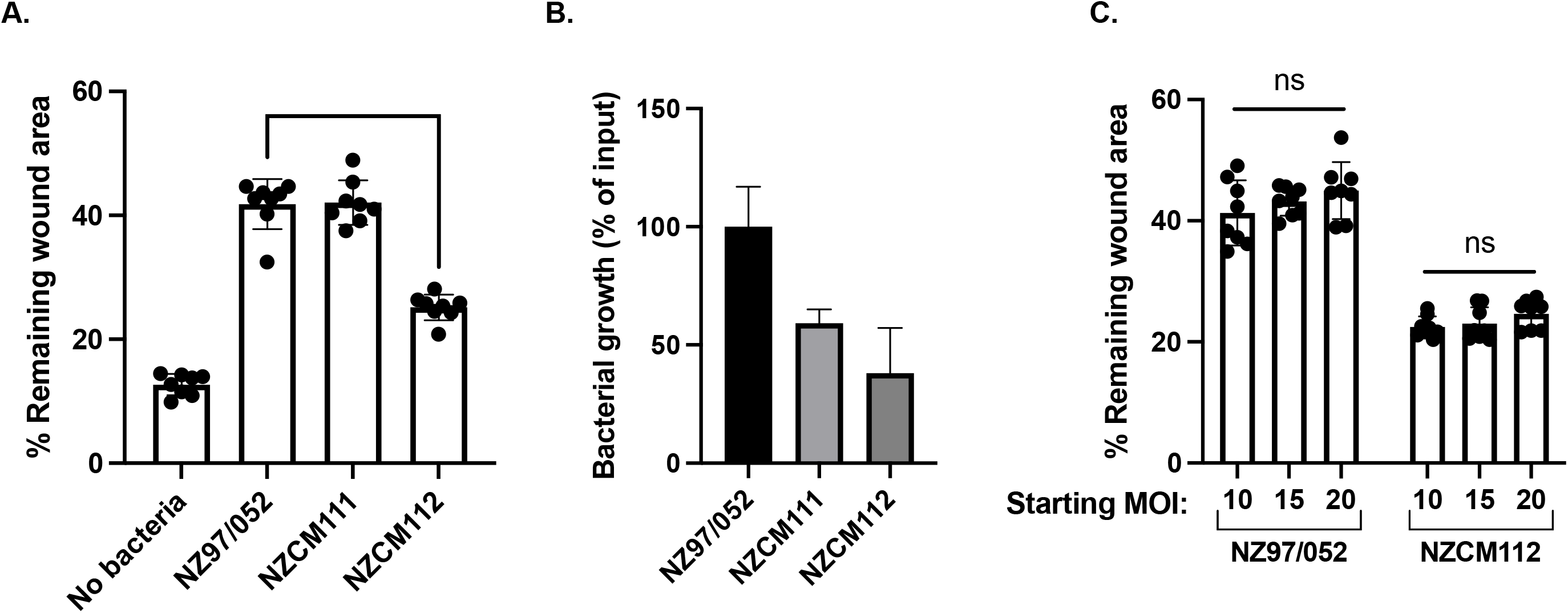
Carriage isolate NZCM112 lacks the ability to inhibit wound closure in vitro. **(A**) The household contact isolate group varies in their ability to inhibit wound repair. The disease-associated isolate, NZ97/052, and carriage-associated isolate, NZCM111, inhibit wound repair, but the carriage-associated isolate, NZCM112, is impaired. (**B**) The growth of NZCM112 is slightly impaired relative to NZCM111 and NZ97/052 after 16 hours in the presence of epithelial cells. (**C**) For isolates NZ97/052 and NZCM112, the starting multiplicity of infection (MOI) does not alter wound repair inhibition after 16 hours. Significance was calculated using an unpaired t test. **** indicates a p value <0.0001, ns, not significant.

### Whole genome sequencing identified household isolate variations

Although NZ97/052, NZCM111 and NZCM112 were indistinguishable by laboratory typing and collected from individuals from a single household, clear phenotypic differences between the isolates were detected, including growth and ability to inhibit wound repair. Our previous studies have identified phenotypic and genomic differences between related isolates from household carriage groups (38). . *N. meningitidis* is highly genetically variable due to recombination between related strains, random mutagenesis, and phase variation. To identify the genetic differences underpinning the different phenotypes, we carried out whole genome sequence analysis of NZ97/052, NZCM111 and NZCM112. Genomic DNA was purified from the three *N. meningitidis* isolates, and sequenced with Illumina MiSeq, using paired-end sequencing. Reads of NZ97/052, NZCM111, and NZCM112 were aligned against the *N. meningitidis* NMBG2136 reference genome (RefSeq genome NC-017513.1), a serogroup B isolate belonging to ST-8 clonal complex (39). The resulting genome sequences were examined to identify where NZCM112 differed from NZ97/052 and NZCM111. Variable haplotypes were examined to ensure that changes within a codon were annotated correctly. A total of 49 haplotypes were identified, with 17 of these found only in NZCM112, but not NZ97/052 or NZCM111. Of the 17 variants that occurred only in NZCM112, 12 were outside coding regions, with 8 in upstream sequence (<100 bp from a predicted start codon) and 4 in intergenic sequence (>100 bp from a predicted start codon). Of the remaining 5 variants in coding sequences, two were predicted missense mutations and two were predicted synonymous mutations, neither of which were predicted to have a major impact on protein phenotype. The remaining variant was the replacement of a six-base sequence, CGGGGG, with a five-base sequence, CAGCA. This mutation occurred in the gene encoding HpuA, a haptoglobin-hemoglobin uptake protein (40, 41). The mutation, resulting in a frameshift and premature stop codon in *hpuA*, was found only in the carriage isolate NZCM112. The *hpuA* variant was the only one of the 49 total variants that was predicted to result in the loss of expression of a protein, and this loss was only seen in NZCM112, but not NZ97/052 or NZCM111. Therefore, *hpuA* was identified as a top candidate gene.

While other frameshift variants were found in the NZ97/052 household, these occurred in a single gene (NMBG2136_1563) which encodes a putative deoxyribonuclease. However, these were not considered as candidates, as the variants occurred in both NZ97/052 and NZCM112 and could not account for the difference in wound repair inhibition between these isolates. The four frameshift mutations effectively cancelled each other out and did not result in a premature stop codon. When the NZ97/052 and NZCM111 alleles of NMBG2136_1563 were entered into the PubMLST allele database, they were found to be identical to alleles in unrelated *N. meningitidis* isolates, suggesting a possible crossover event with another carriage isolate (42).

Although *hpuA* has previously been described as a phase-variable gene in *N. meningitidis*, the mutation we found occurred just upstream of the homopolymeric G tract (43). However, the frameshift resulted in a premature stop codon, occurring just after the homopolymeric G tract, resulting in a predicted 23-amino acid truncated protein. The *hpuA* gene is organized in an operon just upstream of the gene encoding its functional partner, *hpuB*, and both are co-transcribed from a single promoter (41, 43). Previous studies have shown that loss of *hpuA* transcription, due to a frameshift and premature stop codon, has a polar effect, resulting in loss of *hpuB* expression (41). Therefore, *hpuB* was also considered a candidate gene.

### Deletion of *hpuA* reduces inhibition of host cell wound repair

The genome sequence data revealed that an intact HpuA protein was not predicted to be synthesized in NZCM112, while previous studies have shown that when the *hpuA* gene is in the phase-variable “off” configuration, *hpuB* expression is lost or reduced (41). As the *hpuA* gene was shown to be intact in NZ97/052, our aim was to generate an allelic replacement of both *hpuA* and *hpuB*. The operon organization of these genes made it a challenge to ensure that only *hpuA* was targeted, particularly since complementation of genetic mutations in *N. meningitidis* is not easily done via plasmid. We therefore constructed three allelic replacements in the NZ97/052 parental background: two single mutants in *hpuA* and *hpuB* alone, and a double *hpuA-hpuB* mutant. While disruption of *hpuA* is likely to have a polar effect on the expression of *hpuB,* the disruption of *hpuB*, the downstream gene, is not predicted to alter *hpuA* expression (43). All three mutants were generated through the insertion of an antibiotic resistance cassette into the targeted coding region. We then tested the mutants to see if they recapitulated the phenotype of NZCM112.

The three mutants were first compared to the wild type in an in vitro wound repair assay. As seen in Fig 2A, the disruption of *hpuA* (in both the single *hpuA* deletion and the double *hpuA-hpuB* deletion) abrogated the ability of *N. meningitidis* to inhibit host cell migration. There was no defect in wound repair inhibition for the *hpuB* mutation. These results suggest that HpuA, but not HpuB, is the key meningococcal factor for the inhibition of wound repair.

**Figure 2.**
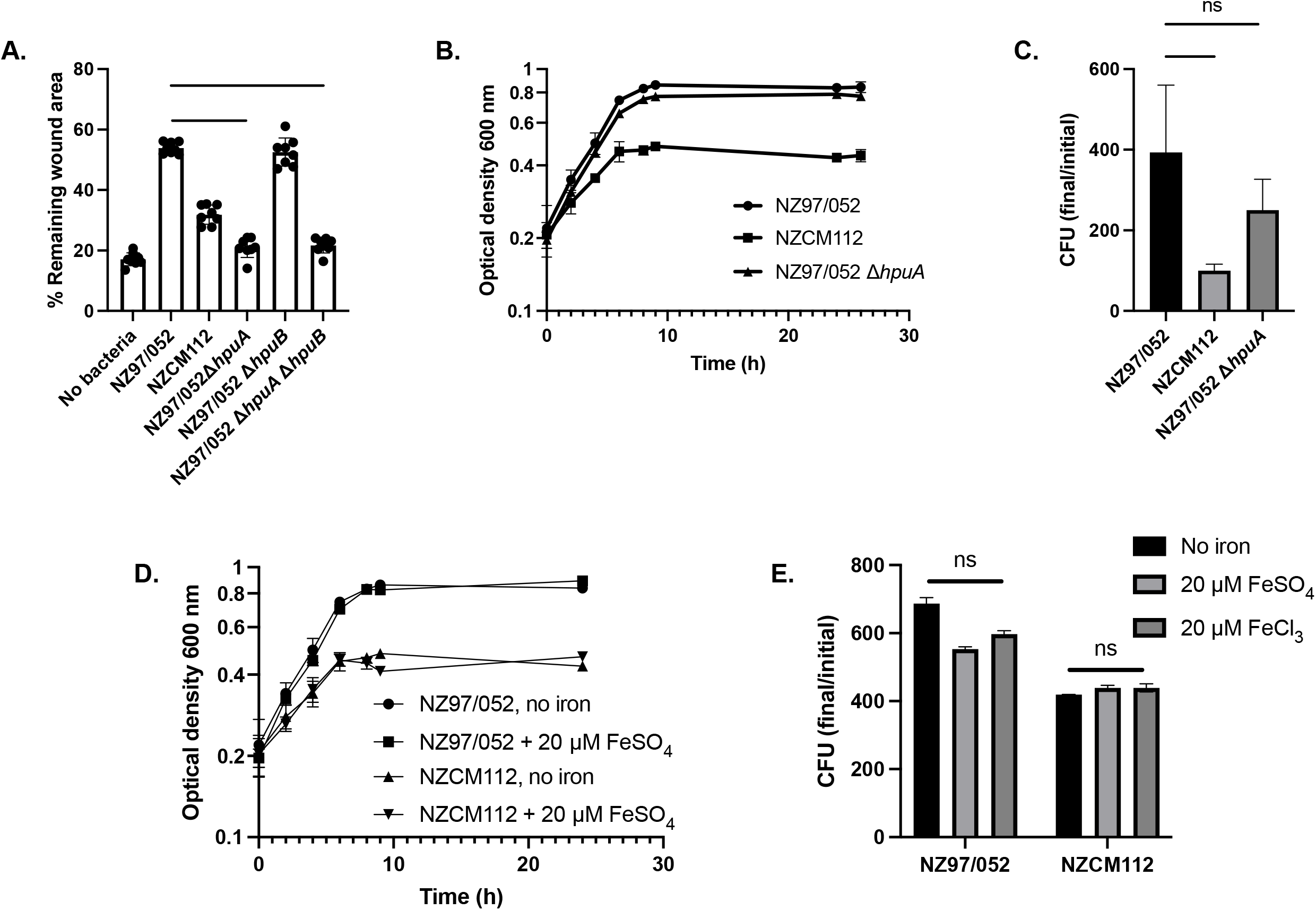
An intact *hpuA*, but not *hpuB,* gene is required for inhibition of host cell wound repair. **(A**) The ability to inhibit wound repair was abrogated when the *hpuA* gene was deleted via allele replacement. This was seen in both the *hpuA* mutant and the *hpuA hpuB* double mutant. The *hpuB* mutant, by contrast, was not impaired for inhibition of wound repair. (**B**) The growth of NZCM112 in standing liquid medium is impaired relative to NZ97/052, but NZ97/052 Δ*hpuA* has only a minor growth impairment. NZCM112 reaches stationary phase at a lower cell density. (**C**) Growth of the isolates in the presence of host epithelial cells was assessed at the endpoint of the wound repair experiment (16 hrs). The growth impairment of CM112 relative to NZ97/052 is significant, but there is no significant difference between NZ97/052 and the Δ*hpuA* mutant. (**D**) Supplementation of the cultures with iron (20 µM FeSO_4_) did not affect the growth rate or the final cell density at stationary phase, for either NZ97/052 or NZCM112. (**E**) Endpoint enumeration of bacteria in the presence of epithelial cells in the presence of FeSO_4_ or FeCl_3_ revealed that iron supplementation did not affect the growth of either NZ97/052 or NZCM112, as assessed after 16 hours of growth. Significance was calculated using an unpaired t test for 2A, 2C, and 2E. **** indicates a p value <0.0001, * indicates a p value <0.05, ns, not significant.

We also asked if the deletion of *hpuA* resulted in a growth defect similar to that seen for NZCM112. As shown in Fig 2B, the *hpuA* mutant had a slight growth defect, compared to the wild type parent, when grown in liquid culture over a time course; however, it did not have the same defect as NZCM112. These results suggest that the additional SNP variants seen in NZCM112 cause this isolate to grow more slowly during exponential phase, and to reach stationary phase at a lower cellular concentration. The basis of this reduced growth remains unknown but is likely due to additional factors than just the loss of *hpuA*. Because HpuA is known to play a role in iron acquisition from the host, we also assessed growth in the presence of epithelial cells, under conditions that mimicked those of the wound repair assay. The bacterial strains were grown in co-culture with epithelial cells and the number of viable colonies was assessed at the end point of 16 hours. As with the liquid culture, the growth of NZCM112 was impaired relative to either the NZ97/052 wild type or the isogenic *hpuA* mutant (Fig 2C). While the Δ*hpuA* mutant may have had a slight growth impairment relative to NZ97/052, this did not reach statistical significance.

These data suggested that the growth defect of NZCM112 was not due to the loss of HpuA, but rather was the result of other SNPs in the genome. However, because of the role HpuA plays in the acquisition of iron for the bacteria from the host, we asked if iron supplementation could rescue the growth defect of NZCM112. Growth was measured by OD_600_ in liquid standing culture over a time course (Fig 2D) and in co-culture with epithelial cells at an endpoint via CFU (Fig 2E). Both strains, NZ97/052 and NZCM112, reached slightly higher cell density when the culture was supplemented with 20 µM FeSO_4_, but the growth of NZCM112 was not rescued by iron supplementation in either case. As seen in Fig 2E, similar results were seen with both Fe^2+^ (FeSO_4_) and Fe^3+^ (FeCl_3_).

Together these data demonstrate that the *N. meningitidis* iron acquisition protein HpuA plays a role in the inhibition of wound repair in epithelial cells, and that this effect is not due to impaired growth of the isolate in the conditions tested. The unexpected role of HpuA in inhibition of host cell migration led us to formulate and test several hypotheses for how this could be occurring.

### *N. meningitidis* HpuA does not alter epithelial cell migration in a haptoglobin-dependent manner

The *N. meningitidis* bipartite receptor components HpuA and HpuB have been shown to efficiently extract heme from hemoglobin and haptoglobin-hemoglobin complexes to provide iron for the bacteria (40, 43–45). HpuB is an outer membrane protein, while HpuA is an acylated lipoprotein partner that interacts with HpuB to bind hemoglobin, haptoglobin-hemoglobin complex, or apo-haptoglobin (41, 45). While one study found that HpuA did not, in the absence of HpuB, detectably bind hemoglobin, haptoglobin-hemoglobin complex, or apo-haptoglobin, HpuA is known to be critical for optimal recognition and use of ligands (46).

The majority of the iron in the human body is stored in hemoglobin and ferritin complexes, and free iron is quickly sequestered by transferrin or lactoferrin. About two thirds of iron in the human body stored in hemoglobin in erythrocytes. If it is released following lysis, haptoglobin binds with high affinity, protecting the body from the oxidative properties of heme and facilitating its removal by macrophages (47, 48). In addition to its role as an acute phase blood protein, haptoglobin is recognized to have a role as an antioxidant and to play a role in the response to infection (49). Haptoglobin has also been shown to have bacteriostatic properties, particularly against Gram-negative pathogens, and to be induced by microbial antigens, such as LPS (49, 50). It has also been shown to be expressed in airway epithelial cells, and in one model system, haptoglobin was shown to increase the rate of cellular migration, possibly by providing iron to migrating cells (49, 51). Because early studies demonstrated binding of HpuAB to apo-haptoglobin (40), we first investigated whether these interactions could play a role in the inhibition of wound repair, independently of the role of HpuAB in heme acquisition.

To test whether the bacteria were binding or sequestering haptoglobin from host cells, exogenous human haptoglobin was added to the migrating cells in the presence of the *N. meningitidis* strains. As seen in Fig 3A, the addition of haptoglobin did not significantly alter the rate of wound repair, even in the absence of bacteria. This is in contrast to previous reports that haptoglobin enhances wound repair, although this could be due to the different types of cells tested (51).

**Figure 3.**
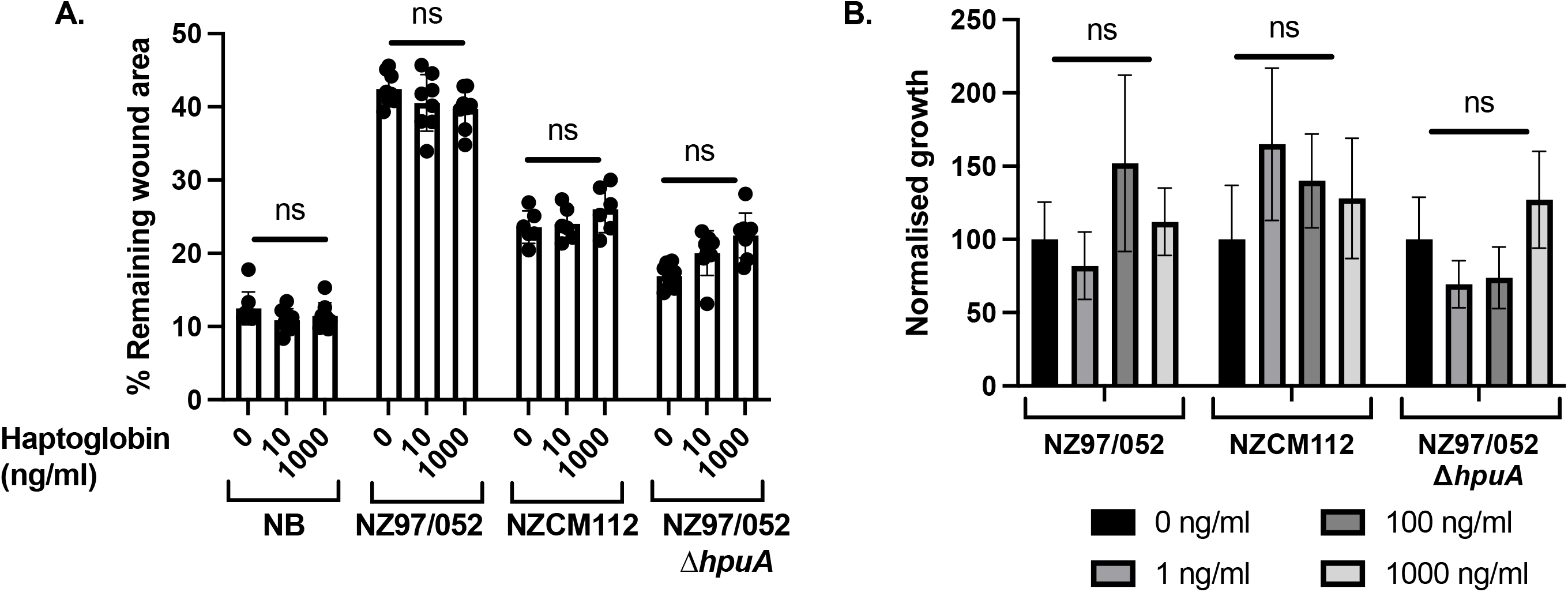
Exogenous human haptoglobin does not influence cellular migration and does not exhibit a bacteriostatic effect on *N. meningitidis*. **(A**) Haptoglobin has been reported elsewhere to influence the rate of cellular migration. When purified human haptoglobin was added to the *N. meningitidis* co-cultures, however, we did not see any significant changes in the rate of wound repair. No effect was seen in the absence of bacteria either, in contrast to other reports. (**B**) Haptoglobin has been reported to have bacteriostatic activity against microbes, particularly Gram-negative bacteria. In a growth assay in the presence of haptoglobin, however, no bacteriostatic effect was detected. Significance was calculated using an unpaired t test. ns, not significant; NB, no bacteria added.

Recent studies have identified a role for haptoglobin in innate immune defenses. Specifically, it has been shown to be bacteriostatic against the Gram-negative pathogens *Proteus mirabilis* and *Pseudomonas aeruginosa* (50). *N. meningitidis*, along with many other human-adapted pathogens, uses haptoglobin-hemoglobin complexes as an iron source, and antimicrobial activity of haptoglobin against these bacterial species has not, to our knowledge, been reported. To test whether haptoglobin has an antimicrobial effect on *N. meningitidis* in the presence of host cells, meningococci were incubated in the presence of confluent 16HBE cells at an approximate MOI of 10 (50). At the same time, purified human haptoglobin (ranging from 1 ng/ml to 1000 ng/ml) was added to the co-cultures. After 24 hours, viable bacteria were determined by CFU counts. In contrast to previous reports, where the highest haptoglobin concentration (1000 ng/ml) reduced bacterial growth for *P. mirabilis* and *P. aeruginosa* to 50% compared to samples without haptoglobin, inhibition of growth was not seen for any of the meningococcal isolates tested (Fig. 3B). The meningococcal strains tested were all resistant to the bacteriostatic activity of haptoglobin, suggesting that factors other than HpuA mediate the innate resistance of *N. meningitidis* to haptoglobin. Thus the bacteriostatic potential of haptoglobin is unlikely to explain the reduced inhibition of cell migration for NZCM112 or NZ97/052 Δ*hpuA*.

### HpuA alters interactions between *N. meningitidis* and host epithelial cells

We did not discover any role for HpuA in enhancing bacterial growth through iron acquisition, protecting the bacteria from the reported antimicrobial effects of haptoglobin, or. sequestering haptoglobin in a way that affects wound repair. We therefore next investigated the interactions of the wild type NZ97/052 parent and the Δ*hpuA* mutant with host cells. We carried out fluorescence microscopy of the isolates, NZ97/052, NZCM112, and NZ97/052 Δ*hpuA*, in co-culture with 16HBE bronchial epithelial cells. The co-cultures were stained with DAPI to image both host cell nuclei and adherent bacteria (Fig 4A-C). The wild type parent strain, NZ97/052, appeared to form clusters of microcolonies that adhered to the epithelial cells (Fig 4A, arrows). The carriage isolate, NZCM112, by contrast, did not form any identifiable microcolonies and appeared to adhere to the epithelial cells at a low rate (Fig 4B). The isogenic *hpuA* deletion mutant in the NZ97/052 parental background similarly appeared to adhere in much smaller numbers, and microcolonies appeared to be fewer and less dense (Fig. 4C). We investigated the adherence of these three strains to 16HBE cells using a standard assay to quantify cell association. Confluent 16HBE cells were co-cultured with the different *N. meningitidis* isolates for one hour, after which cells were washed thoroughly to remove all non-adherent bacteria, and then lysed with saponin, which does not affect the viability of meningococci. CFUs representing total cell-associated bacteria (including both adherent and intracellular) were enumerated. The cell association of each strain was calculated as a percentage of the initial inoculant, also measured by CFU enumeration (Fig 4D). The NZ97/052 isolate displayed about 4.6-fold higher levels of association with 16HBE cells relative to NZCM112 (p < 0.0001). The NZ97/052 Δ*hpuA* isolate significantly decreased association with 16HBE cells relative to the wild type parent (p < 0.001). The numbers of NZ97/052 CFU associated with 16HBE cells was approximately 1.9-fold higher relative to the Δ*hpuA* mutant.

**Figure 4.**
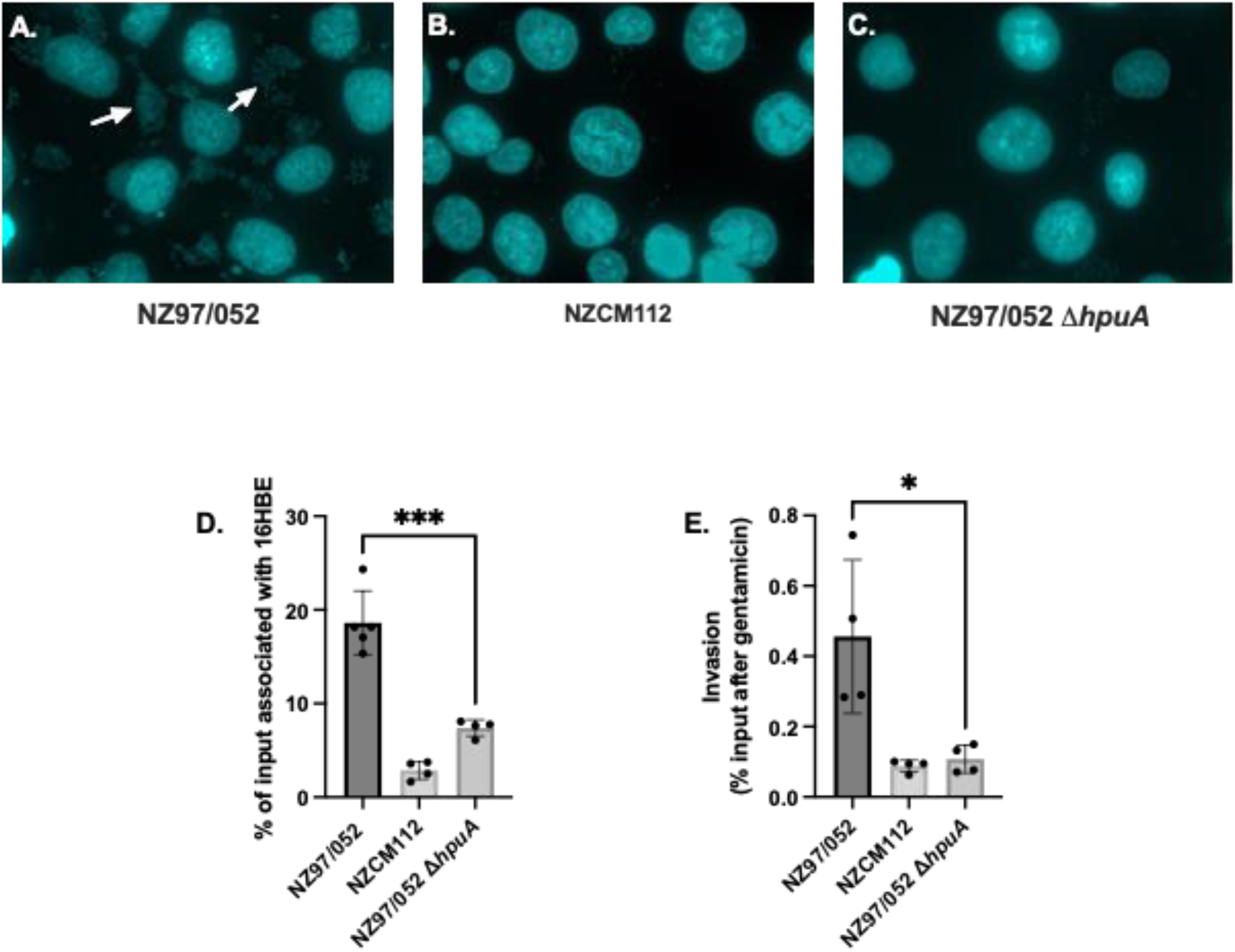
*N. meningitidis* strains lacking HpuA have reduced adherence and invasion of bronchial epithelial cells. Fluorescence microscopy of the bacteria co-cultured with 16HBE cells was carried out with DAPI staining to visualize the adherent bacteria and host cell nuclei. (**A**) NZ97/052 adheres in high numbers to 16HBE cells, forming adherent clusters (microcolonies; arrows). (**B**) The carriage isolate NZCM112 adheres in much lower, nearly undetectable, numbers. (**C**) The isogenic Δ*hpuA* mutant of NZ97/052 also apparently has a reduced ability to adhere to epithelial cells. (**D**) Quantification of cell-association (including both adherent and intracellular bacteria) after 1 hour of co-culture confirms that both NZCM112 and NZ97/052 Δ*hpuA* are associated with epithelial cells at significantly lower numbers. (**E**) A gentamicin protection assay to quantify intracellular bacteria after 4 hours of co-culture reveals that a reduced ability to invade epithelial cells correlates with reduced adherence and cell association. Significance was calculated using an unpaired t test. *** indicates a p value <0.001, * indicates a p value <0.05.

*N. meningitidis* has also been shown to invade epithelial cells, possibly as a means of accessing nutrients or evading the immune response. We therefore carried out gentamicin protection assays to determine whether the increased bacterial association led to increased cellular invasion. The assay was carried out similarly to the cell association assay, but the bacteria and cells were co-incubated for a longer period (4 hours, rather than 1 hour), after which gentamicin was added to kill any extracellular bacteria. The NZ97/052 isolate displayed significantly higher levels of invasion of 16HBE cells than NZCM112 (Fig 4E; p < 0.05), an approximately 2.3-fold increase in intracellular bacteria. The NZ97/052 isolate had an approximately 4.2-fold higher rate of invasion into 16HBE cells, relative to NZ97/052 Δ*hpuA*, a significant difference (p < 0.05).

These results support our initial observation that HpuA plays a role in the interaction between the disease-associated NZ97/052 isolate and host epithelial cells. Tight adherence is a prerequisite for intracellular invasion, which may occur in mucosal epithelial cells as a means of escape from the local immune response. The microscopy also demonstrated an interesting pattern of adherence of NZ97/052 to host cells; it appeared that NZ97/052 formed many microcolonies on the surface of the host cells, a configuration not seen with NZCM112 or NZ97/05 Δ*hpuA*. Together this work identifies a previously unrecognized role for HpuA, as a bacterial adhesin for epithelial cells.

### Heterologous expression of HpuA confers an adherent phenotype

We wished to restore the *hpuA* gene to either NZCM112 or the NZ97/052 Δ*hpuA* mutant, to see if the adherent phenotype could be restored. However, we were unable to complement the *hpuA* mutation by restoring it on a plasmid, due to a lack of access to replicating plasmids for *N. meningitidis*. Instead, we opted to express HpuA in *E. coli* BL21(DE3), which lacks a native *hpuA* homologue. A synthetic *E. coli* codon-optimized DNA fragment of *hpuA* was designed, with the slipped-strand mispairing error-prone phase variable homopolymeric G-tract removed (43, 52). The HpuA protein was expressed in *E. coli* when induced with IPTG, but not with 1% glucose to repress protein expression (Fig 5A). We have previously observed that *E. coli* strains are toxic to bronchial epithelial cells, so we were unable to do any experiments that involved a lengthy co-incubation (e.g., wound repair inhibition or gentamicin protection assay). During initial experiments, the 16HBE cells were monitored via microscopy to ensure that they retained integrity. The expression of HpuA was induced with IPTG for one hour, as this was found to result in sufficient expression of HpuA with no observable *E. coli* growth defect (Fig 5A). The uninduced control was treated with 1% glucose, to further repress HpuA expression. Both induced and uninduced *E. coli* cultures were added to confluent 16HBE cells for 30 minutes and non-adherent bacteria were removed by thorough washing and lysis with saponin. The cell-associated bacteria were calculated as a percentage of the initial inoculant, both measured by CFU enumeration (Fig 5B). The average association of *E. coli* not expressing HpuA was 0.40% of the input bacteria after 30 minutes. By contrast, 0.83% of the bacteria expressing HpuA were adherent, a significant difference (p < 0.01). This represents an approximately 2-fold increase in association with respiratory epithelial cells when HpuA was expressed.

**Figure 5.**
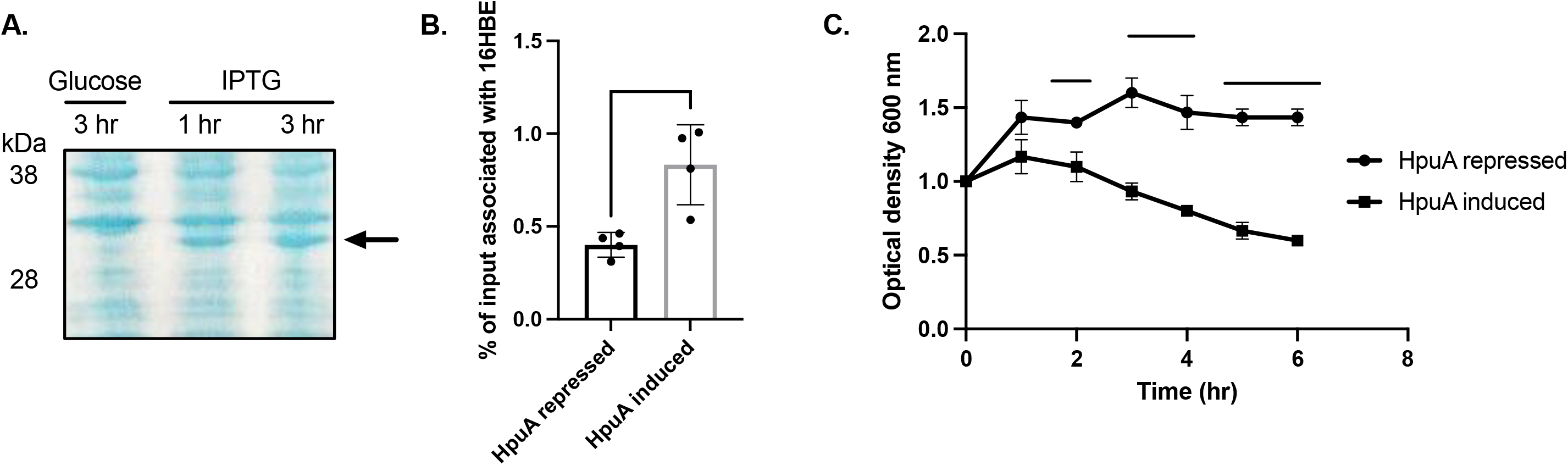
Heterologous expression of HpuA in *E. coli* BL-21 cells confers adherent and autoaggregative phenotype. (**A**) A synthetic codon-optimized version of HpuA was expressed in *E. coli* BL-21 cells. When induced with 0.4 mM IPTG, a protein close to the predicted size of HpuA (35 kDa) was detected (arrow). The protein was not made under repressive conditions (1% glucose). (**B**) When the HpuA protein had been induced for one hour, BL-21 cells were added to 16HBE cells for 30 minutes, after which the cells were washed, lysed with saponin, and CFUs enumerated relative to input. BL-21 cells expressing HpuA were cell-associated at about twice the rate of cells not expressing HpuA. (**C**) To investigate the potential effect of HpuA on aggregation, BL-21 cells were grown and HpuA induced with IPTG. The cultures were then adjusted to an OD_600_ of 1 and left undisturbed for 6 hours. A small sample of the bacterial suspension was carefully taken from the top of the sample periodically and the OD_600_ calculated. A reduction of the OD_600_ at the surface was interpreted as aggregation of the bacteria. Significance of cell association was calculated using an unpaired t test. Significance of sedimentation was calculated for individual time points. **** indicates a p value <0.0001, *** indicates a p value of <0.001, ** indicates a p value <0.01.

The clusters of NZ97/052 bacteria adherent to epithelial cells, which was not seen with the Δ*hpuA* mutant or the carriage CM112 strain (Fig 4A-C), led us to hypothesize that HpuA could be playing a role in bacterial aggregation. We assayed the ability of HpuA expression to confer an aggregative phenotype on the host *E. coli* strain, which normally does not auto-aggregate. Sedimentation profiles (Fig 5C) showed that the sedimentation rate of the strain expressing HpuA was significantly greater than the non-expressing *E. coli* control (p < 0.0001 at hours 5 and 6). This assay suggests that HpuA is a major determinant of the aggregation of *N. meningitidis*, although other proteins with redundant function have been reported (53, 54).

These data together demonstrate that HpuA has at least two previously unknown roles, in mediating adherence to host epithelial cells and aggregation. Cumulatively, these functions enable the bacteria to adhere better to host cells and to form microcolonies that may promote colonization and persistence in the airways.

### Iron supplementation alters the interactions between epithelial cells and *N. meningitidis* strains that lack HpuA

While carrying out these experiments, we asked if HpuA could be impacting epithelial cells indirectly as a result of key differences in the metabolism of carriage- and disease-associated isolates. Specifically, we hypothesized that isolates that inhibit wound repair could be more readily consuming a nutrient, such as iron, needed by the host cells for migration. This could result in host cell deprivation of iron, required for multiple enzymes that play a role in wound repair and cellular migration.

To test this hypothesis, we supplemented our co-cultures of epithelial cells with *N. meningitidis* isolates with iron (20 µM FeSO_4_) and observed whether this altered the inhibition of wound repair. If bacteria were consuming iron, and depriving host cells of it, supplementation of the co-cultures with iron would have been expected to increase cell migration (and reduce the inhibition of wound repair) by NZ97/052. However, we found that iron supplementation did not alter the ability of NZ97/052 to inhibit wound repair, but that NZCM112 and NZ97/052 Δ*hpuA* both gained the ability to inhibit wound repair (Fig 6A). Wound repair inhibition remained unchanged for NZ97/052, and the overall rate of wound closure in the absence of bacteria did not change with iron supplementation. Instead, the addition of iron appeared to switch the NZCM112 and NZ97/052 Δ*hpuA* strains to a phenotype that resembled that of NZ97/052. The limited effect of the iron on cells in the absence of added bacteria suggested that the supplemented iron is altering the meningococcal strains that lack HpuA, not the host cells.

**Figure 6.**
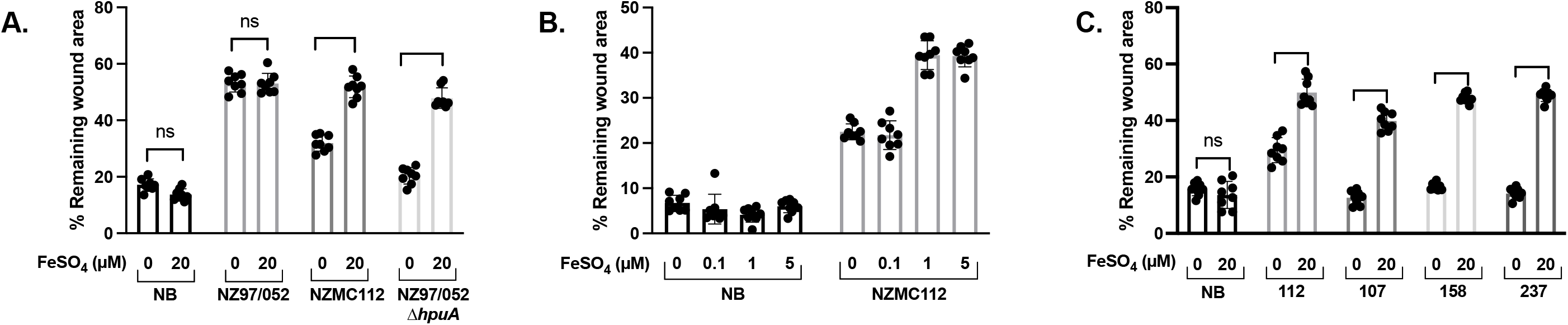
Iron supplementation confers the ability of strains lacking HpuA to inhibit wound repair. (**A**) The addition of 20 µM FeSO_4_ to the in vitro wound repair assay did not alter the ability of NZ97/052 to inhibit wound repair, nor did it alter the migration of cells without any added bacteria. However, supplementation with iron conferred the ability to inhibit wound repair on both NZCM112 and NZ97/052 Δ*hpuA*, with the wounds inhibited to a similar degree as seen with the wild type NZ97/052. (**B**) Varying concentrations of FeSO_4_ were added, revealing that the altered phenotype occurred with 1 µM FeSO_4_ added, but not with 0.1 µM. (**C**) Additional unrelated carriage isolates, previously shown to not be able to inhibit wound repair (35), acquired the ability to inhibit wound repair under conditions of iron supplementation. Carriage strains tested were NZCM112, NZCM107, NZCM158, and NZCM237 and are indicated in the graph by their last three numbers. Significance was calculated via pairwise t-tests. *** indicates a p value <0.001, **** indicates a p value of <0.0001, ns, not significant. NB, no bacteria added.

We tested a range of concentrations of FeSO_4_ supplementation to determine if there was a dose-dependent effect. However, as seen in Fig 6B, it did not appear to be dose-dependent, as no effect was seen with 0.1 µM FeSO_4_ added, while similar effects were seen with supplementation with 1 µM and 5 µM FeSO_4_. This indicates more of an “on-off” transition mechanism, rather than a dose-dependent response.

We asked whether this phenomenon was limited to our closely related isolates, or if it could be observed in other, unrelated, meningococcal isolates that do not inhibit wound repair. A range of unrelated carriage isolates (obtained from nasopharyngeal swabs from healthy volunteers) that we had previously shown not be able to inhibit wound repair, were tested with and without supplementation with 20 µM FeSO_4_ (35). These strains belong to different sequence types and a range of serogroups (including serogroups B, C, and non-groupable). As seen in Fig 6C, the same phenomenon was observed for each of these strains, where the addition of iron to the co-culture medium did not alter the migration of uninfected epithelial cells but conferred the ability to inhibit wound repair on the carriage isolates.

Because our previous results had shown that the inhibition of host cell wound repair correlated with adherence to the cells, we next investigated whether the addition of iron altered the numbers of NZCM112 and NZ97/052 Δ*hpuA* cells associated with epithelial cells, as a percentage of the input bacteria (FIG 7A). The association of NZ97/052 with epithelial cells did not change with the addition of iron to the cell culture medium. However, the addition of iron increased the association of NZCM112 with host cells, although it was not restored to similar levels as seen with NZ97/052. While we noted an increase in cell association for the NZ97/052 Δ*hpuA* deletion strain, this did not reach statistical significance (p = 0.08). A similar effect was seen when we investigated cellular invasion, using a gentamicin protection assay. As seen in Figure 7B, intracellular bacteria, as a percentage of input bacteria, increased significantly for both NZCM112 and NZ97/052 Δ*hpuA* when iron was added to the medium, with no change seen for NZ97/052. Despite the significant increase, the total percentage of intracellular bacteria did not reach the level of those seen for the NZ97/052 wild type.

**Figure 7.**
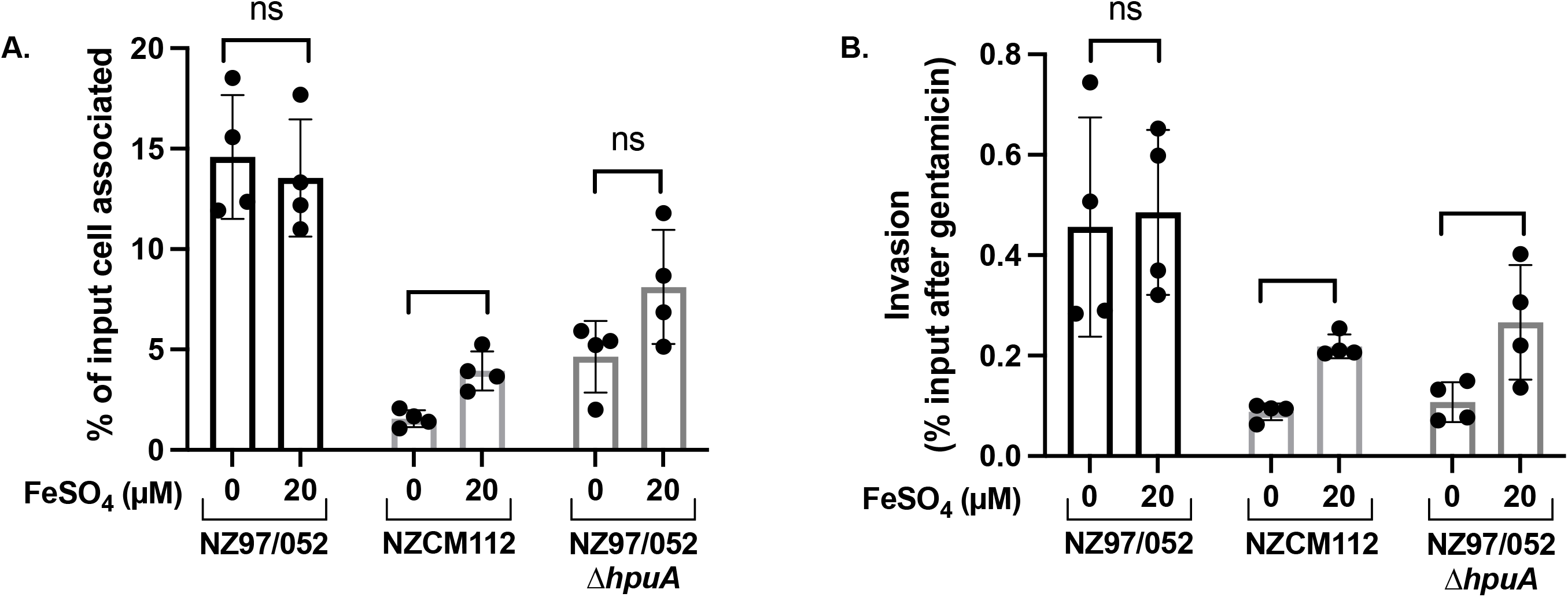
Iron supplementation enhances the cell association and invasion of strains lacking HpuA. (**A**) The addition of 20 µM FeSO_4_ did not alter the degree of association of NZ97/052 with 16HBE cells, but it did increase the association of NZCM112 and NZ97/052 Δ*hpuA*. The increase was significant for NZCM112, but did not reach significance for NZ97/052 Δ*hpuA* (p=0.08). (**B**) Intracellular meningoocci were enumerated after four hours of co-culture. Iron supplementation (20 µM FeSO_4_) did not alter cellular invasion by NZ97/052, though it increased invasion for both NZCM112 and NZ97/052 Δ*hpuA*. Significance was calculated via pairwise t-test. * indicates a p value <0.05, ** indicates a p value <0.01, and *** indicates a p value >0.001, ns, not significant.

## Discussion

Multiple moonlighting functions of the meningococcal iron-acquisition lipoprotein, HpuA, are described here. We compared a set of naturally-occurring household contact isolates for their ability to inhibit host cell wound repair, using an in vitro model of re-epithelialisation, an early step in wound repair, identifying a household contact group that differed in their ability to inhibit wound repair. Whole genome sequencing of this set of isolates suggested a role for the HpuAB receptor in host cell interactions. Construction of mutants confirmed a role for HpuA, but not HpuB. We subsequently demonstrated that HpuA mediates adherence, invasion, and inhibition of wound repair of epithelial cells, as well as bacterial aggregation.

Nutritional immunity, the withholding of key nutrients by the host, has long been known as an important aspect of resistance to colonizing bacteria (55). The bipartite HpuAB receptor plays a key role in host iron piracy by *N. meningitidis*. HpuA is a surface-exposed lipoprotein, while HpuB is a transmembrane protein; they are encoded by co-transcribed genes previously shown to be regulated by the global iron regulator, Fur (41). *N. meningitidis* strains frequently express a second hemoglobin receptor, HmbR (56, 57). Both HpuAB and HmbR bind hemoglobin, extracting the heme moiety complexed with iron and transporting it into the cell, where the iron is removed and the heme metabolised (44). However, only HpuAB is capable of extracting heme from hemoglobin when it is complexed with haptoglobin (45). A recent structural study of HpuA homologues from *N. gonorrhoeae* and *Kingella denitrificans* revealed a direct interaction betwen HpuA and heme, with conserved residues required for the interaction identified (58). In addition, conserved residues that were dispensable for heme binding were also identified, consistent with our observation that HpuA is a moonlighting protein with other functions.

HpuAB receptors are found in both pathogenic and non-pathogenic *Neisseria*, while HmbR is associated with pathogenic isolates (59). Most *Neisseria* strains have at least one of the HpuAB or HmbR receptors, though strains with only HpuAB are more highly associated with carriage (59). However, more than 90% of isolates from certain hypervirulent clonal complexes were found to express both HpuAB and HmbR (60). In one case, the phase variable *hpuA-hpuB* locus was found to have been in the “off” phase in the inoculating isolate, but in the “on” phase following accidental human passage (61). HmbR is more strongly associated with invasive disease in *N. meningitidis* and has been shown to be essential for infection in an infant rat model (56, 62, 63). HmbR is not expressed, however, in *N. gonorrhoeae*, as the sequence contains a premature stop codon, leaving the gonococcus reliant on expression of HpuAB (57). Indeed, the *hpuA-hpuB* locus has been shown to be expressed by *N. gonorroheae in vivo* in the female genital tract early in infection, suggesting a role for it in the mucosa (64).

Multiple environmental factors that enhance damage to the airways have been shown to increase the risk of developing invasive meningococcal disease, suggesting that this is one way meningococci access deeper tissues (30–34). HpuA, by inhibiting wound closure, may be important for this characteristic of meningococci, although the mechanisms by which HpuA inhibits epithelial cell wound repair remain unknown. Our previous study found that inhibition of wound repair correlated with actin morphology, including the formation of actin cables and lamellipodia (35). Other bacterial factors that have been shown to inhibit wound repair, such as the *Pseudomonas* Type-III secreted effectors ExoST, and the *Helicobacter pylori* VacA cytotoxin, specifically inactivate host cell cytoskeletal structural or signaling proteins, inhibiting host cell migration (65–68). However, specific interactions between HpuA and the host cytoskeleton have not been identified.

We also investigated a role for haptoglobin in altered wound repair, as early reports demonstrated direct binding interactions between HpuAB and hemoglobin, haptoglobin-hemoglobin, and apo-haptoglobin (40). Haptoglobin has recently been shown to be synthesized in airway epithelial cells, where it is induced by bacterial lipopolysaccharide; it has also been reported as having a bacteriostatic effect on some Gram-negative pathogens, and as promoting wound repair in some types of cells (49–51, 69). However, we did not find any evidence that HpuA is involved in evading the antimicrobial action of haptoglobin, or in the sequestration of haptoglobin. Specifically, the addition of exogenous haptoglobin did not alter cell migration for any of the samples, nor were any of our meningococcal strains susceptible to the reported anti-microbial effects of haptoglobin. This is consistent with more recent reports that have suggested that there is no direct interaction between HpuA and haptoglobin (58).

Our studies of the interactions between meningococci, with or without HpuA, and bronchial epithelial cells revealed that HpuA is a major adhesin for these cells, at least with our clinical isolates. Microscopy of co-cultures revealed differences in adherence and the formation of microcolonies in strains lacking HpuA. Specifically, deletion of HpuA reduced adherence of the bacteria to host cells to about 40% of the wild type level. Heterologous expression of HpuA in *E. coli* conferred an adherent phenotype onto these bacteria. We also found that HpuA promotes the invasion of epithelial cells by *N. meningitidis*. Host cell invasion by *N. meningitidis* has been reported as occurring with both hypervirulent and carriage-associated strains, and may represent a means of evading aspects of the host immune response or antibiotics (4, 70–72). These results were surprising, as type IV pili have long been shown to be a major adhesin for meningococci with epithelial cells (73, 74). The isolates in our study were piliated, though we have previously shown that inhibition of wound repair is independent of Type IV pili (35). Despite the dominant role of pili in the adherence of meningococci to epithelial cells, alternative adhesins have been reported, including the opacity proteins, Opa and Opc, which mediate intimate adhesion, and autotransporters App, NadA, and NhhA (75–82). Non-pathogenic *N. cinerea* has been reported to bind to epithelial cells in a Type IV pilin-independent manner, although the adhesin was not identified (23, 83). The observation that strains lacking HpuA were less likely to form microcolonies led us to identify an additional moonlighting function for HpuA, in mediating aggregation. Aggregation may provide bacteria with protection against the host immune response or antimicrobial agents, and can abet escape from phagocytosis (84, 85). Meningococcal aggregation likely plays a role in the formation of microcolonies on the surface of host cells, important for bacterial colonization and persistence (53, 72). The meningococcal Type-IV pili and trimeric autotransporter AutA have been shown to mediate aggregation in *N. meningitidis* (53, 54). Both aggregation and adherence to host cells are critical functions for meningococci, and are mediated by redundant factors.

When *N. meningitidis* is in the presence of epithelial cells and a layer of mucus, the bacteria express low levels of classical adhesins, but high levels of iron transporters, a finding consistent with the scarcity of iron in mucus (11, 12). In this system, small numbers of bacteria were still observed in close association with host cells, independent of Type IV pili (11). This close association may enable the bacteria to access intracellular iron sources (15). Meningococci accelerate ferritin degradation in epithelial cells and consume a breakdown product of the ferritin complex as an iron source (15). This process begins when the bacteria are extracellular but attached to epithelial cells as microcolonies, as well as when the bacteria are intracellular. HpuA, by enabling adherence of meningococci to host cells, may contribute to iron piracy by both acquiring heme from hemoglobin, as well as by enabling adherent bacteria to access iron from the host cell.

One of our more surprising findings was that supplementation of iron to co-cultures of *N. meningitidis* with host cells altered the ability of strains lacking HpuA (e.g., NZCM112 or the Δ*hpuA* mutant) to adhere, invade, and inhibit wound repair. The addition of iron to these strains made them behave more like the HpuA-expressing strains. Supplementation with iron did not alter wound repair of epithelial cells in the absence of any bacteria, suggesting that the iron was affecting the bacteria, rather than the host cells directly. We had hypothesized that iron limitation, resulting from bacterial iron piracy, could have impacted the ability of host cells to migrate. Although human hemoglobin is not abundant in cell culture, the HpuAB receptor has been shown to be able to acquire heme from bovine sources, present in the fetal calf serum, which we reasoned might be important for wound repair (57). However, the evidence did not support this hypothesis, as the addition of iron would have been predicted to increase wound repair in the presence of potentially all of the bacterial strains, but particularly those strains expressing HpuA, and this was not observed. Furthermore, iron supplementation did not result in any appreciable difference in bacterial growth rates, suggesting the results were not due to altered bacterial growth. One possible explanation is that iron supplementation induced the expression of an additional meningococcal protein that also plays a role in the inhibition of wound repair. The effect of this additional unknown protein could be masked in cells that already express HpuA, and only apparent in strains lacking HpuA. Alternatively, iron supplementation could result in repression of HpuAB expression in a Fur-dependent manner, while iron-induced expression of this second protein compensates for reduced levels of HpuA (43). A study of *N. meningitidis* genes induced following the addition of iron revealed at least 153 genes upregulated in the presence of iron, including 49 hypothetical genes encoding proteins of unknown function (86). Of genes encoding proteins of known function, most were categorized as having roles in energy metabolism, protein synthesis, and cell envelope assembly. Other meningococcal proteins known to be upregulated by iron include metal exporters, which reduce the cytotoxic effects of excess metals. While metal exporters are normally situated in the inner membrane, they may interact with unknown outer membrane porins to expel toxic levels of transition metals (87). Another iron-inducible protein that could mediate adhesion is the TonB-dependent transporter ZnuD (TdfJ in *N. gonorrhoeae*) (16, 88). Our results suggest there are additional meningococcal factors, including possible additional moonlighting proteins, that mediate adhesion and inhibition of wound repair, still remaining to be identified.

Moonlighting bacterial proteins are increasingly recognised for their important role in pathogen virulence (89). Moonlighting enables pathogen economy, where a single protein can fulfill two or more crucial functions for life in the mucosa while limiting the number of potential immune antigens displayed to the host. Moonlighting also contributes to redundancy of essential functions, which enables phase variation in meningococci. A previously identified moonlighting meningococcal protein is fructose-1, 6-bisphosphate aldolase, which is localized to both the cytoplasm and the outer membrane (90). Meningococcal aldolase also mediates adherence to human brain microvascular endothelial cells and binds human plasminogen via its C-terminal lysine residue (90, 91). Another meningococcal metabolic enzyme that localizes onto the cell surface is the metabolic enzyme glyceraldehyde 3-phosphate dehydrogenase (GAPDH), although no additional moonlighting function was noted (92). Additional cytosolic enzymes have been reported as being located on the outer membrane and binding plasminogen, highlighting the ubiquity of this pathogenic strategy for *N. meningitidis* (93). Moonlighting functions of iron-acquisition proteins have not been noted for *N. meningitidis*, but they have for other bacterial residents of iron-poor mucosal surfaces, including *Streptococcus* species and *Haemophilus influenzae* (94, 95). The widespread nature of heme acquisition protein moonlighting has particularly been noted in *H. influenzae*, which, while unrelated to *N. meningitidis*, occupies a similar host niche (95). Moonlighting functions of *H. influenzae* heme acquisition transporters include adherence to host proteins, import of glutathione, resistance to antimicrobial peptides, and acquisition of potassium (95). Multi-functional virulence proteins in bacteria are likely much more widespread than is currently appreciated, and additional moonlighting functions of iron, heme, and zinc transporters of pathogenic *Neisseriae* are likely yet to be identified, offering a rich area of future investigation.

## Material and Methods

### Bacterial strains and growth conditions

*N. meningitidis* were collected as described during a household contact study carried out in Auckland in the late 1990s (36). Isolates were maintained by the Meningococcal Reference Laboratory (MRL) at the Institute of Environmental Science and Research (ESR), as part of routine surveillance of meningococcal disease in New Zealand. The carriage and disease meningococcal isolates in this study are described in Table 1. They were collected, analyzed by laboratory typing, and maintained as described previously (38). *N. meningitidis* was grown on Columbia Blood Agar (CBA) plates (Fort Richard Laboratories, Auckland), for routine passage, or on BBL Brain Heart Infusion agar (Oxoid) plates, supplemented with kanamycin (50 µg/ml; Fluka, Switzerland) where required, at 36°C in a humidified 5% CO_2_ incubator. *Escherichia coli* BL21(DE3) Hi-Control (Lucigen) was used as an expression host for heterologous expression of HpuA. *E. coli* was grown on Luria-Bertani agar plates or in LB broth, supplemented with kanamycin (50 µg/ml).

**Table 1.** Meningococcal isolates used in this study.

| Isolate | Source | Typing (serogroup: serotype: serosubtype; sequence type) |
| --- | --- | --- |
| NZ97/052 | Blood sample | C:2b:P1.2,5; ST-8 |
| NZCM111 | Nasopharyngeal swab | C:2b:P1.2,5; ST-8 |
| NZCM112 | Nasopharyngeal swab | C:2b:P1.2,5; ST-8 |
| NZCM107 | Nasopharyngeal swab | B:nt:19-15; ST-178 |
| NZCM158 | Nasopharyngeal swab | NG:1:P1.22,14-6,36.2;ST-2154 |
| NZCM237 | Nasopharyngeal swab | NG:nt:P1.16; unknown ST |
| NZ97/052 $\Delta hpuA$ | Constructed by gene deletion | C:2b:P1.2,5; ST-8 |
| NZ97/052 $\Delta hpuB$ | Constructed by gene deletion | C:2b:P1.2,5; ST-8 |
| NZ97/052 $\Delta hpuAB$ | Constructed by gene deletion | C:2b:P1.2,5; ST-8 |

NG, nongroupable; nt, nontypable; ST, Sequence type. All original isolates were collected as part of a household contact carriage study (36). Gene deletions were constructed as described in the methods.

Three of the *N. meningitidis* isolates used in this study were derived from a single household. The NZ97/052 disease-associated isolate was derived from a patient with meningococcal disease in 1997; the NZCM111 and NZCM112 isolates were derived from the same household but were isolated from healthy carrier household contacts of the patient. All three household isolates were identified as belonging to the strain type C:2b:P1.2,5 and the clonal complex ST-8. The other NZCM isolates (NZCM107, NZCM158, and NZCM237) were collected during the same household contact study, but from different households, and were described previously (35, 36).

### Meningococcal growth assays

The growth of meningococcal isolates was measured over a time course by optical density or at an end point by enumeration of colony forming units (CFU). For the time course, meningococci grown overnight on CBA plates were collected with a sterile swab and resuspended in BHI broth (Oxoid). The bacterial suspension was adjusted to OD_600_ = 0.2 and 15 ml was added to an empty T-75 flask (Jet Biofil, China), along with nutritional supplementation as required (i.e., FeSO_4_ or FeCl_3_). Flasks were incubated flat in the incubator, to provide aeration for the liquid culture. Every hour, cultures were mixed by pipetting up and down and 50 µl of the culture was removed to measure optical density. Measurements were taken in triplicate and the experiment was performed three separate times.

Growth of meningococcal isolates was also measured by enumeration of colony forming units under conditions designed to mimic those of the Oris cell migration assay. Meningococci grown overnight on CBA plates were resuspended in M199 medium (Gibco, USA) with 10% inactivated FCS. Dilutions of the bacterial suspension were plated out on CBA plates to quantify the inoculant. The resuspended bacteria were then added at an approximate MOI of 10 to confluent 16HBE cultures that had been cultured in 24-well plates and serum-starved for 24 hours prior to addition of the bacteria. Plates were sealed with a Breathe-Easy membrane (Sigma-Aldrich) to prevent internal contamination. After 16 hours of incubation, the cell culture supernatant was diluted and plated out on CBA. Isolates were assayed in parallel with three to five replicates per isolate. Experiments were performed three independent times.

### Whole genome sequencing

Genomic DNA was purified from meningococci with the Gentra Puregene Yeast/Bacteria kit (Qiagen) according to the manufacturer’s instructions, with the following changes. Bacteria, grown on CBA plates overnight, were scraped from the plate (∼10 µl), resuspended in 300 µl of lysis buffer, and incubated at 56°C for 1 hour to kill meningococci, after which the remaining steps were followed. DNA concentration and quality were assessed by electrophoresis, 260/280 ratio, and the Quant-iT PicoGreen double-stranded DNA (dsDNA) assay kit (Thermo Fisher). Isolate genomes were sequenced by Genomics Aotearoa (Dunedin, New Zealand). Samples were prepared using the TruSeq Nano library format and sequenced by Illumina MiSeq using paired-end sequencing (Illumina, USA). Adaptor trimmed reads were quality trimmed using Trimmomatic-0.32 at a Phred quality greater than 20 and a minimum length of 100 bp (96). Paired reads were aligned to RefSeq genome NC_017513.1 (NMBG2136) using bowtie 2.1.0 (39). NMBG2136, an ST-8, serogroup B isolate, was selected as a reference genome since belongs to the same sequence type as the NZ97/052 household, although they differ in serogroup. Nucleotide variants, single nucleotide variants (SNPs) and small insertions and deletions were identified using haplotype-ware Freebayes 1.0 and annotated using SnpEff (97, 98). Where there were numerous variants in a gene sequence, recombination was investigated by entering the allele sequences into the PubMLST allele database to see which species it was associated with (42).

### Construction of *N. meningitidis* mutants

Meningococcal mutants were made by allelic replacement of the targeted gene with an AphA3 kanamycin (Km) resistance cassette by natural transformation (99). *N. meningitidis* NZ97/052 genomic DNA was used as a template to amplify about 200 bp of flanking regions of each target gene (*hpuA* and/or *hpuB*). Primers were designed based on the published sequence of NMBG2136 and are listed in Table 2 (39). All internal primers include a long sequence of overlap with the Km primers. The AphA3 Km cassette in pUC19 (pUC19 Km) was used as a template for a third PCR, using the primers Km_Fw_1 and Km_Rv_1. For each mutagenic construct, the three PCR products were verified and combined to set up a fusion PCR, amplifying with the outermost primers. The resulting fusion PCR product was verified, further amplified and concentrated, then used directly for a natural transformation of NZ97/052. To generate a *hpuA-hpuB* double mutant, sequences upstream of *hpuA* and downstream of *hpuB* were combined with the Km PCR product and used for natural transformation of NZ97/052. Natural transformations of *N. meningitidis* were performed using purified, concentrated PCR products, as described previously (35). The correct insertion of the Km cassette into the genome was confirmed by PCR and Sanger sequence analysis of the targeted gene region.

**Table 2.**
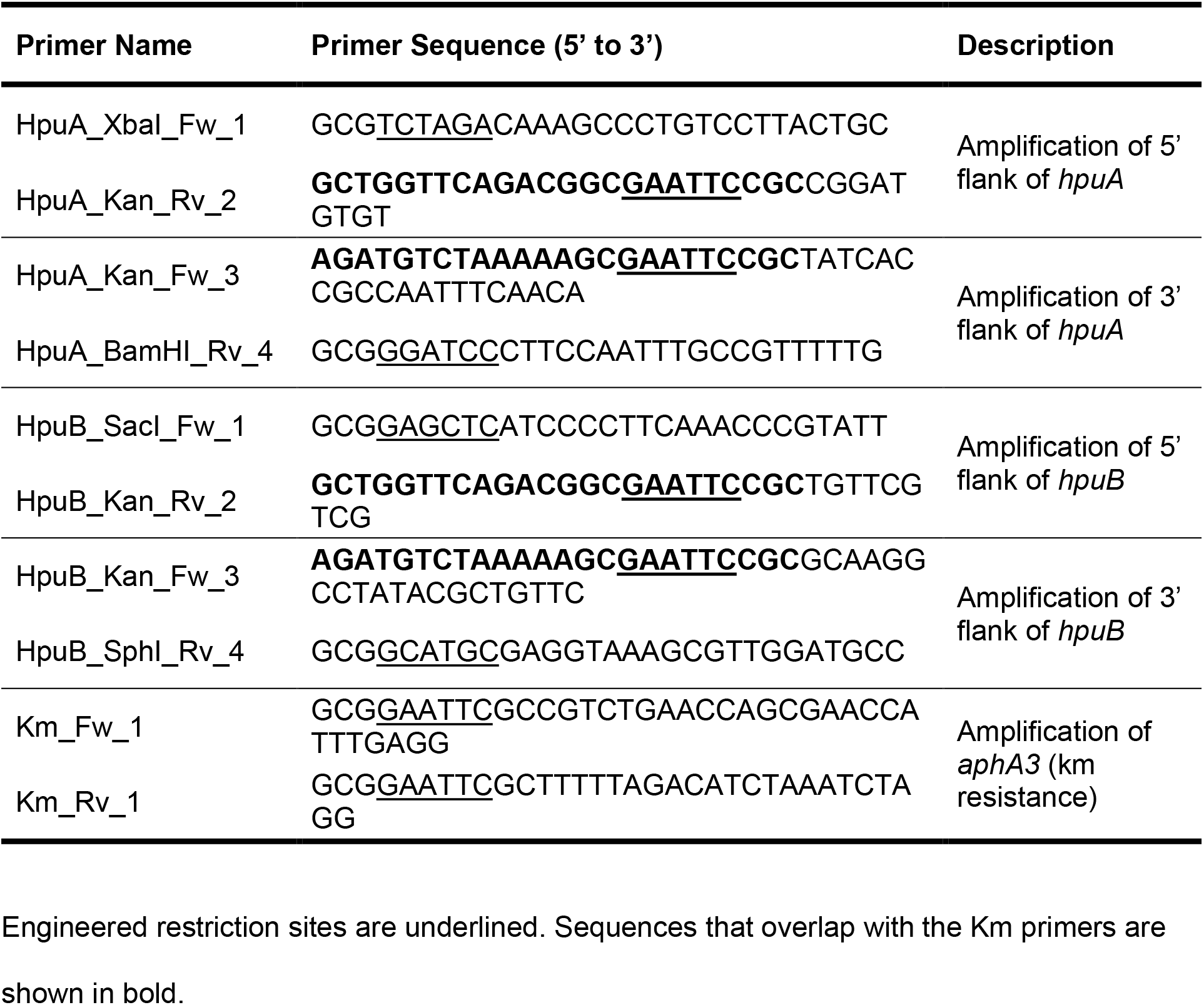
Primers used in this study.

### Heterologous expression of *hpuA* in *E. coli*

Because we were unable to complement the NZ97/052 Δ*hpuA* strain, we opted to heterologously express HpuA in an *E. coli* strain, which lacks an endogenous *hpuA* gene. The *hpuA* gene sequence from NMBG2136 (NMBG2136_1863) was downloaded from NCBI (https://www.ncbi.nlm.nih.gov/gene/12411497). The sequence was adjusted to improve expression and stability, optimizing codons for *E. coli* using the Codon Optimization Tool (Integrated DNA Technologies) and removing the homopolymeric G-tract to reduce the likelihood of a slipped-strand mispairing error. Restriction sites (NdeI and XhoI) were added to the gene, with the ATG start codon incorporated into the engineered NdeI site, and the stop codon removed before the XhoI site. The codon-optomized synthetic gene was synthesized and cloned into the NdeI and XhoI sites of the pET28a(+) vector (Novagen) by Twist Biosciences. The synthetic plasmid, containing a His-tagged version of HpuA and a linker sequence at the N-terminus, was resuspended in TE and introduced into BL21(DE3) Hi-Control chemically competent cells (Lucigen), which were selected on LB plates supplemented with kanamycin.

### Induction of HpuA in *E. coli*

One colony containing the synthetic plasmid was grown in 5 ml of LB broth supplemented with kanamycin at 37°C with shaking, until the culture reached an OD_600_ of 0.6. This starter culture was split in two tubes. HpuA expression was induced in one tube by the addition of 0.4 mM IPTG and repressed in the other by the addition of 1% glucose. The growth rate of induced and uninduced cultures were compared to screen for potential toxicity. Following induction, the total cell protein fraction was collected for induced and repressed HpuA-expressing *E. coli* at 1 and 3 hours. Optical density was used to normalize cell loading volumes of samples. Cell pellets were resuspended in 100 µl PBS, then 100 µl of 2X sodium dodecyl sulfate (SDS) sample buffer (Thermo Fisher Scientific) was added, the sample was passed through a 27-guage needle several times, heated for 3 minutes at 85°C, and stored at -20°C until gel analysis. Heterologous expression of HpuA was confirmed by protein gel analysis (Novex 10-20% Tris-Glycine mini wedge well gels, Thermo Fisher Scientific) followed by Coomassie blue staining (Bio-Safe Coomassie G-250, Bio-Rad, USA).

### Cell culture and infection

Bronchial respiratory epithelial cells (16HBE14o-, abbreviated as 16HBE) were routinely cultured in M199 medium supplemented with 10% inactivated fetal calf serum (FCS) (100). For infection experiments, 16HBE cells were suspended at 6 x 10^4^ cells/ml and cultured in 6- or 24-well plates until confluent. For cell migration assays, 16HBE cells were suspended at a density of 3 x 10^5^ cells/ml in M199 (Gibco) plus 10% FCS, with 100 µl of cell suspension added to each well of a collagen-coated Oris cell migration assay 96-well plate (Platypus Technologies, Madison WI), which was incubated overnight to allow the cells to adhere. All wells in an Oris cell migration plate contain a stopper insert to prevent the cells from adhering to the center of the well. The Oris cell migration assay with 16HBE cells and meningococcal infection was carried out and analyzed essentially as previously described (35). For immunofluorescence microscopy, 16HBE cells were grown on 8-well glass slides (Nunc Lab-Tek II chamber slide system; Sigma-Aldrich) with each well seeded with 450 µl of cell suspension at 1.2 x 10^5^ cells/ml and grown for 24 hours, until confluent. Where noted, cell cultures were supplemented with iron or haptoglobin at various concentrations. Supplements were made by combining crystals of ferrous sulfate heptahydrate (Fe^2+;^ Toronto Research Chemicals, Sapphire Biosciences, Australia) or ferric chloride hexahydrate (Fe^3+^; Sigma-Aldrich) with distilled water to a 1 M concentration and sterilizing with a 0.22 µm filter. Each solution was diluted as needed and made fresh for each experiment. To assess the impact of haptoglobin on cell migration, haptoglobin from pooled human plasma (H3536, Sigma-Aldrich) was resuspended in cell culture medium to the required concentrations, then added to the Oris migration assay.

### Cell association and invasion assays

Confluent 24-well plates of 16HBE cells were serum-starved for 24 hours, then washed and medium with serum was added. Cells were infected with a fresh suspension of *N. meningitidis* for a final MOI of approximately 10. Immediately after adding the bacteria to the wells, 10 µl samples were taken for dilution and enumeration of CFUs. Cells and bacteria were co-incubated for one hour for cell association assays, and four hours for gentamicin protection assays. After incubation, cells were washed with warm PBS three times to remove all bacteria that were not adherent or intracellular. Cells were lysed with 500 µl 1% saponin (Sigma-Aldrich) in PBS per well and incubated at room temperature for 10 minutes with gentle shaking. Dilutions of the cell lysate were made and plated onto CBA plates.

To determine the number of intracellular meningococci, the same method was used, but prior to the addition of saponin, 500 µl of gentamicin (150 µg/ml) (Thermo Fisher Scientific) was added for one hour to kill extracellular bacteria as described (101). Gentamicin was removed and cells were thoroughly washed with PBS to remove all traces of gentamicin before addition of saponin. The cell lysates were diluted and plated onto CBA plates. Following overnight incubation, CFU were counted; input and output numbers of CFU were used to calculate percentage of intracellular or cell-associated bacteria.

Cell association of *E. coli* was determined essentially as for meningococci, with a few alterations. *E. coli* was first grown in liquid culture rather than on plates (LB with 50 µg/ml kanamycin) to mid-log phase, then split into two cultures, with one induced with IPTG, and the other repressed with glucose, for one hour. The OD_600_ was measured, and the *E. coli* added to confluent 16HBE cells at a MOI of approximately 10. The 16HBE cells were only co-cultured with *E. coli* for 30 minutes, to reduce toxicity to the cells. Dilutions of input bacteria were carried out as for meningococci, and 16HBE cells were washed and lysed with saponin, and bacterial CFU enumerated on LB plates supplemented with kanamycin.

For all assays, isolates were tested in parallel with three to five replicates per isolate. Experiments were performed three times and statistical significance was tested using unpaired t-test or one-way ANOVA.

### Fluorescence microscopy of meningococci and cells

16HBE cells were cultured to confluence on eight-well chamber slides, as described above. Fresh *N. meningitidis* were used to infect the cells at an MOI of 10. Immediately, 10 µl from each well was removed and dilutions were plated out on CBA plates to determine the input number of bacteria. After four hours of infection, the medium was removed, and cells were washed three times with cold PBS. The cells were immediately fixed with 4% paraformaldehyde in PBS for 20 minutes at room temperature. The paraformaldehyde solution was removed by washing with PBS. Cells were permeabilized with 0.5% Triton X-100 for 10 minutes, blocked with a solution made up with 10% heat-inactivated FCS, 0.1% Triton X-100 in PBS for 20 minutes, then were actin stained with DAPI (4’,6-diamidino-2-phenylindole, dihydrochloride) nucleic acid stain (Thermo Fisher Scientific, USA) in PBS for 10 minutes, before being washed three times with PBS. Slides were mounted with Prolong Gold Antifade Mountant (Thermo Fisher Scientific USA) and stored in the dark at 4°C until analysis. Chamber slides were analyzed at 100X magnification on an Olympus upright fluorescence microscope (BX-51), and images were recorded with a digital camera attached to the microscope. For each well, eight pictures were taken at random positions. Images were merged and analyzed with ImageJ (102).

### Assay for haptoglobin effect on bacterial viability

The antimicrobial effect of human haptoglobin on *N. meningitidis* isolates was assessed using a previously described method (50). Briefly, 16HBE cells were grown to confluence in a 24-well plate and infected with a fresh *N. meningitidis* suspension at an estimated MOI of 10. At the same time, a range of concentrations of pooled human haptoglobin (Sigma-Aldrich), ranging from 1 to 1000 ng/ml final concentration, were added to the co-cultures. Viable bacteria from the co-cultures were enumerated immediately after the infection (to determine the inoculant) and again after 24 hours.

## Acknowledgments

This work was supported by a New Zealand Lottery Health Translational Research Grant and an ESR-VUW joint PhD Scholarship (to GAS). Grant-in-aid of research support was also provided by the Wellington Medical Research Foundation (WMRF) and the New Zealand Wound Care Society (NZWCS).

